# Prepontine non-giant neurons drive flexible escape behavior in zebrafish

**DOI:** 10.1101/668517

**Authors:** Gregory D. Marquart, Kathryn M. Tabor, Sadie A. Bergeron, Kevin L. Briggman, Harold A. Burgess

**Affiliations:** Division of Developmental Biology, *Eunice Kennedy Shriver* National Institute of Child Health and Human Development, Bethesda, MD, USA; Neuroscience and Cognitive Science Program, University of Maryland, College Park, MD 20742, USA; Department Genes – Circuits – Behavior, Max Planck Institute of Neurobiology, 82152 Martinsried, Germany; Biology Department, West Virginia University, Morgantown, WV 26506; Circuit Dynamics and Connectivity Unit, National Institute of Neurological Disorders and Stroke, Bethesda, MD 20892; Research Center caesar, Max Planck Society, Bonn, Germany

**Keywords:** escape response, startle, sensorimotor integration, B3 recombinase, superior vestibular nucleus, Mauthner neuron, zebrafish

## Abstract

Many species execute ballistic escape reactions to avoid imminent danger. Despite fast reaction times, responses are often highly regulated, reflecting a trade-off between costly motor actions and perceived threat level. However, how sensory cues are integrated within premotor escape circuits remains poorly understood. Here we show that in zebrafish, less precipitous threats elicit a delayed escape, characterized by flexible trajectories, that are driven by a cluster of 38 prepontine neurons that are completely separate from the fast escape pathway. Whereas neurons that initiate rapid escapes receive direct auditory input and drive motor neurons, input and output pathways for delayed escapes are indirect, facilitating integration of cross-modal sensory information. Rapid decision making in the escape system is thus enabled by parallel pathways for ballistic responses and flexible delayed actions.

## Introduction

Escape behaviors are fast defensive responses to threats that are typically driven by short sensorimotor reflex arcs (Bullock, 1984). Some species possess multiple modes of escape, including less powerful responses, characterized by delayed initiation and less vigorous motor activity (Comer et al., 1988; Krasne, 1965; von Reyn et al., 2014). Such delayed escape reactions are frequently produced in response to the same stimuli that drive fast escape responses, but preferentially elicited by weaker cues. There has been little work characterizing circuits that mediate delayed escapes (Bhattacharyya et al., 2017), precluding analysis of neuronal mechanisms that select and coordinate threat responses.

Escape behavior, triggered by abrupt tactile, auditory or visual stimuli, has been studied extensively in teleost fish. Central to the escape circuit are the Mauthner cells, a bilateral pair of giant reticulospinal neurons that trigger explosive C-start maneuvers with a single action potential (Eaton et al., 1981, 1977b; Zottoli, 1977). However, a second class of escape swim has also been described (Burgess and Granato, 2007; Eaton et al., 1977a, 1982, 1984; Koyama et al., 2016). Zebrafish larvae respond to auditory stimuli with kinematically distinct short-latency C-starts (SLCs) and long-latency C-starts (LLCs) (Burgess and Granato, 2007; Issa et al., 2011; Jain et al., 2018). Like delayed escapes in other species, LLCs are less vigorous, more variable and preferentially elicited by weaker stimuli. However, neurons which initiate LLCs have not been described, and it is not known whether LLCs share neuronal pathways with SLCs or why the initiation of LLCs is delayed relative to Mauthner mediated responses.

To resolve these questions, we conducted an unbiased circuit-breaking screen to identify specific neurons that drive delayed escapes in zebrafish. We discovered a bilateral cluster of approximately 20 neurons per side in the prepontine hindbrain that are necessary and sufficient to initiate delayed escapes. Prepontine escape neurons are only active on trials where larvae initiate a delayed escape, but do not project directly to the spinal cord, indicating that they act as premotor neurons. Finally, results from behavioral experiments suggested that delayed escapes provide an opportunity for multi-modal integration. Our data reveals that parallel pathways subserve ballistic and flexible delayed escapes, shedding light on the neuronal architecture that enables rapid behavioral choice.

## Results

We used high-speed video to analyze escapes triggered by acoustic/vibrational stimuli in free-swimming 6 day post-fertilization (dpf) larvae (Fig. 1A). Auditory C-start reactions comprise a fast C-bend (C1), counterbend to the other side, and swim bout. As previously described, the distribution of latencies from stimulus onset to C1 initiation was bi-modally distributed: larvae initiated a short latency C-start (SLC) within 12 ms of the stimulus, or a long latency C-start (LLC) between 16-50 ms after the stimulus (Fig. 1B). We also confirmed that individual larvae executed both SLC and LLC escapes on different trials to stimuli of the same intensity (Fig. 1B). Whereas SLCs were highly stereotyped all-or-nothing responses, LLC responses were kinematically variable, showing a significantly greater co-efficient of variation for all C1 movement parameters (Fig. 1C), and less vigorous, resulting in a small net displacement (Fig. 1D). The relatively long reaction time, high variability and low movement speed of LLCs are features shared with secondary modes of escape in other species (Krasne, 1965; von Reyn et al., 2014; Wine and Krasne, 1972).

**Figure 1.**
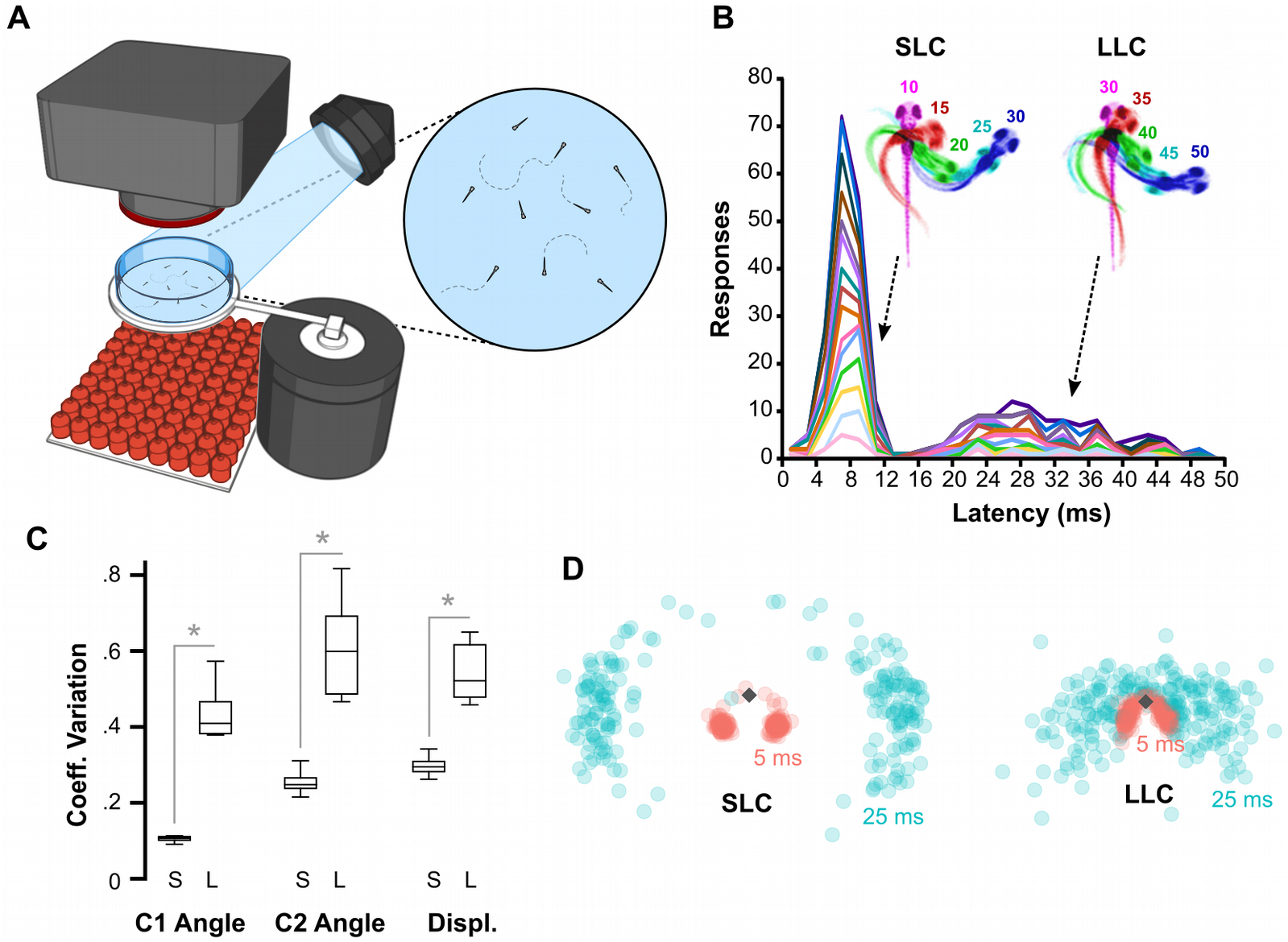
Ballistic and delayed escape reactions performed by larval zebrafish (A) Schematic of behavioral experiments in free-swimming larvae: Groups of 15-20 6 dpf larvae were imaged from above at 1000 frames per second with a high-speed camera. An infrared (IR) LED array below provided illumination. Non-directional acoustic/vibratory stimuli were delivered to the arena by a minishaker. (B) Frequency histogram of response latencies for individual larvae (n=15, color coded). Inset: Timelapse images of initial C-bend for larvae performing an SLC or an LLC, color coded by millisecond post-stimulus. (C) Coefficient of variation (CV) for the initial bend angle (C1), counterbend angle (C2) and net displacement for SLC (S) and LLC (L) responses. n=16 groups of larvae. * p < 0.001. (D) Heatmap of final positions after SLC (n=763 responses) and LLC (n=593 responses) responses.

To identify neurons that subserve delayed escapes, we initiated a circuit-breaking screen using a library of Gal4 lines to selectively ablate subsets of neurons before testing escape behavior (Fig. 2A). We confirmed three lines where LLC responses were reduced by more than 50% after ablation (*y252-Gal4*, *y293-Gal4* and *y330-Gal4*; Fig. 2B-C). Critically, SLC responses were not reduced, excluding impairments in sensory sensitivity, and additional motor phenotypes differed between the lines, presumably due to the distinct sets of neurons ablated in each (Fig. 2D, Fig. S1). We reasoned that the three lines may label a shared population of neurons critical for LLCs, and evaluated overlap in co-registered whole-brain images of Gal4 expression (Marquart et al., 2015). Strikingly, three-way co-localization was restricted to a single area in the prepontine hindbrain: a bilateral region of rhombomere 1 (R1) located dorsolaterally to the locus coeruleus, comprising 19 ± 1.7 neurons per side (mean/s.e.m. for n =10 *y293-Gal4* larvae; Fig. 2E-F, Fig. S2). Neurons in the prepontine cluster were therefore candidates for driving delayed escape behavior.

**Figure 2.**
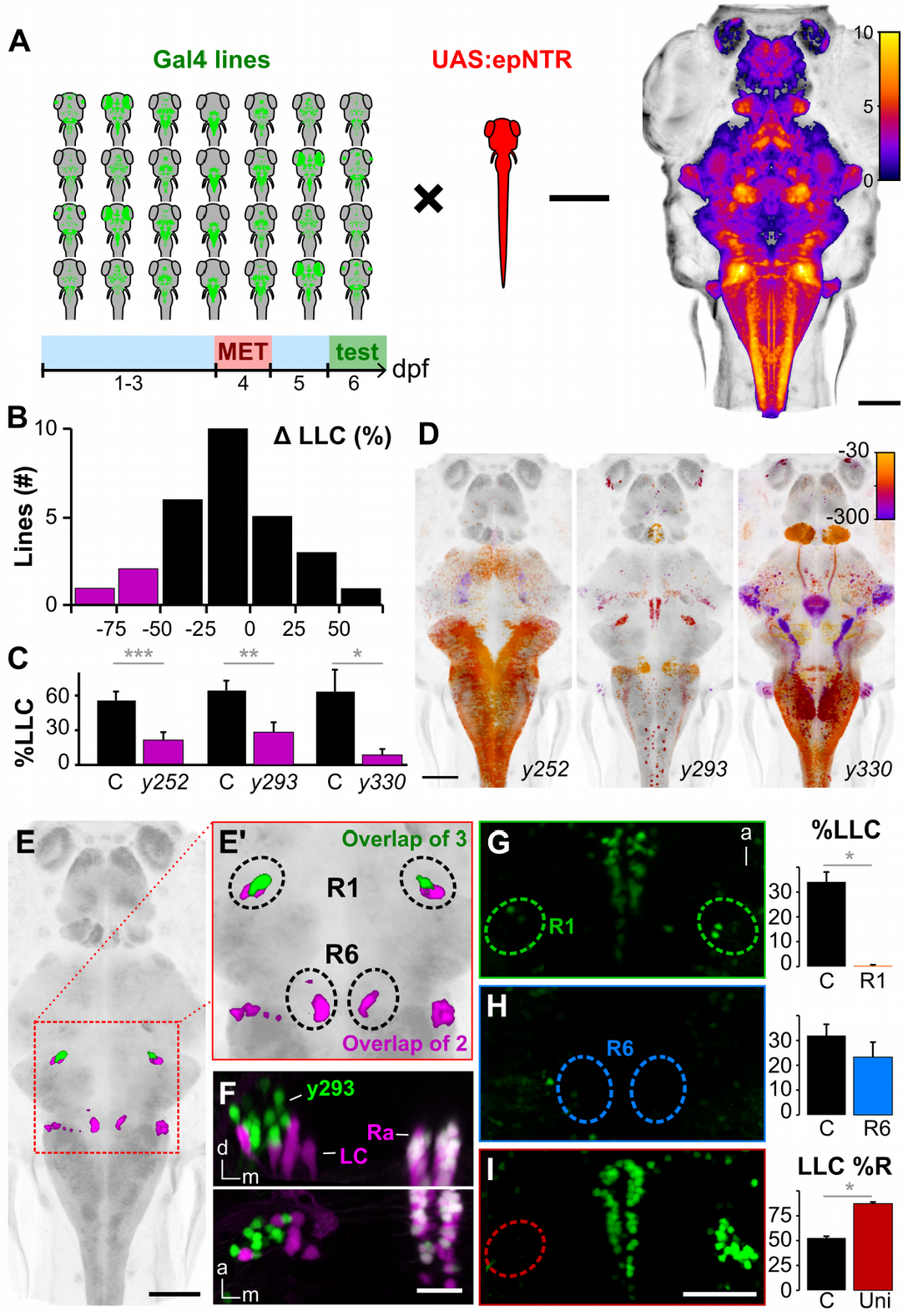
A cluster of neurons in the prepontine hindbrain initiates delayed escapes (A) Schematic of circuit-breaking screen: 28 Gal4 enhancer trap lines were crossed to *UAS:epNTR*, labeled neurons ablated, and tested for escape behavior. Right: heat-map representing brain coverage (number of lines labeling a given voxel). (B) Histogram of the change in LLC probability following ablation (compared to met-treated non-NTR expressing sibling controls) for each line screened. Magenta: Gal4 lines with a >50% reduction. (C) LLC probability for lines highlighted in (B). LLC probability after ablation (magenta) and in met-treated sibling controls (black). *y252* (n=22 control, 31 ablated larvae), *y293* (n=17,17), *y330* (n=8,7). (D) Maximum horizontal projections for Gal4 lines with reduced LLC probability after ablation. Expression is color-coded for depth (μm below image top). (E) Expression overlap between *y252-Gal4* and *y293-Gal4* (magenta) and between all three lines (green). Boxed area enlarged in (E’). (F) Coronal (top) and dorsal (bottom) projections of confocal sub-stacks through the R1 cluster in *y293-Gal4; UAS:Kaede; vmat2:GFP*^*pku2*^ larvae. Arrows indicate locus coeruleus (LC) and raphe (Ra) labeled by *vmat2:GFP*. (G-H) LLC probability after laser ablation of R1 (G, n=9) and R6 (H, n=16) in *y293-Gal4*. * p < 0.05 (I) Percent of LLCs made in a rightward direction after left R1 ablation (Uni, n=14) and non-ablated controls (n=24). * p < 0.05. Scale bars: 100 μm in A, D, E; 40 μm in G-I; 25 μm in F

We next laser ablated prepontine neurons to test if they are required for delayed escapes. Focal ablation of the bilateral clusters completely abolished delayed escapes across all stimulus intensities (Fig. 2G). In contrast LLCs were unimpaired after eliminating a cluster of neurons in R6 that were co-labeled by *y252-Gal4* and *y293-Gal4* (Fig. 2H, Fig. S3A). After unilateral ablation of the prepontine cluster, more than 80% of LLCs were directed toward the intact side (Fig. 2I). Prepontine ablations did not affect motor performance but reduced the probability of fast escapes, an effect that is likely to be non-specific because unilateral lesions did not affect SLC direction (Fig. S3B-D). Taken together, transgenic and laser ablation experiments reveal that a bilateral cluster of neurons in the prepontine hindbrain are essential for delayed escapes. These neurons are adjacent to the locus coeruleus but not labeled by the monoaminergic marker *vmat2* (Fig. 2F). The transgene in *y293-Gal4* is integrated in the first intron of *fibronectin type III domain containing 5b* (*fndc5b*) and therefore likely reflects the spatial expression pattern of this gene (Marquart et al., 2015). In the Allen Mouse Brain Atlas, *Fndc5* is also expressed adjacent to the locus coeruleus, in the vestibular nuclei (Lein et al., 2007; Thompson et al., 2014), a region previously implicated in driving vestibular startle responses in mammals (Fig. S4) (Bisdorff et al., 1994; Li et al., 2001). These and other similarities (see Discussion) suggest that prepontine escape neurons in fish are homologous to the mammalian superior vestibular nucleus.

Prepontine neurons might directly initiate escape reactions, or regulate the responsiveness of another pathway. To test whether prepontine neuron activation is sufficient to drive escape behavior, we expressed the channelrhodopsin variant ChEF in *y293-Gal4* neurons and selectively stimulated prepontine neurons in head-embedded larvae using a digital mirror device (DMD) (Fig. 3A)(Lin et al., 2009). Control ChEF negative larvae did not respond to LED illumination. Unilateral illumination of ChEF positive neurons elicited behavioral responses in 54.6% of trials, of which half were initiated with a large angle tail flexion similar to C-start responses in free swimming fish (Fig. 3B-C, Video S1). C-starts triggered by unilateral optogenetic activation were primarily initiated to the ipsilateral side (76.6% of responses ipsilateral, one-sample t-test versus 50% p = 0.003; Fig. 3D). These results confirm that prepontine neurons drive delayed escapes, and further support the idea that neurons in each hemisphere predominantly, although not exclusively, drive ipsilateral responses.

**Figure 3.**
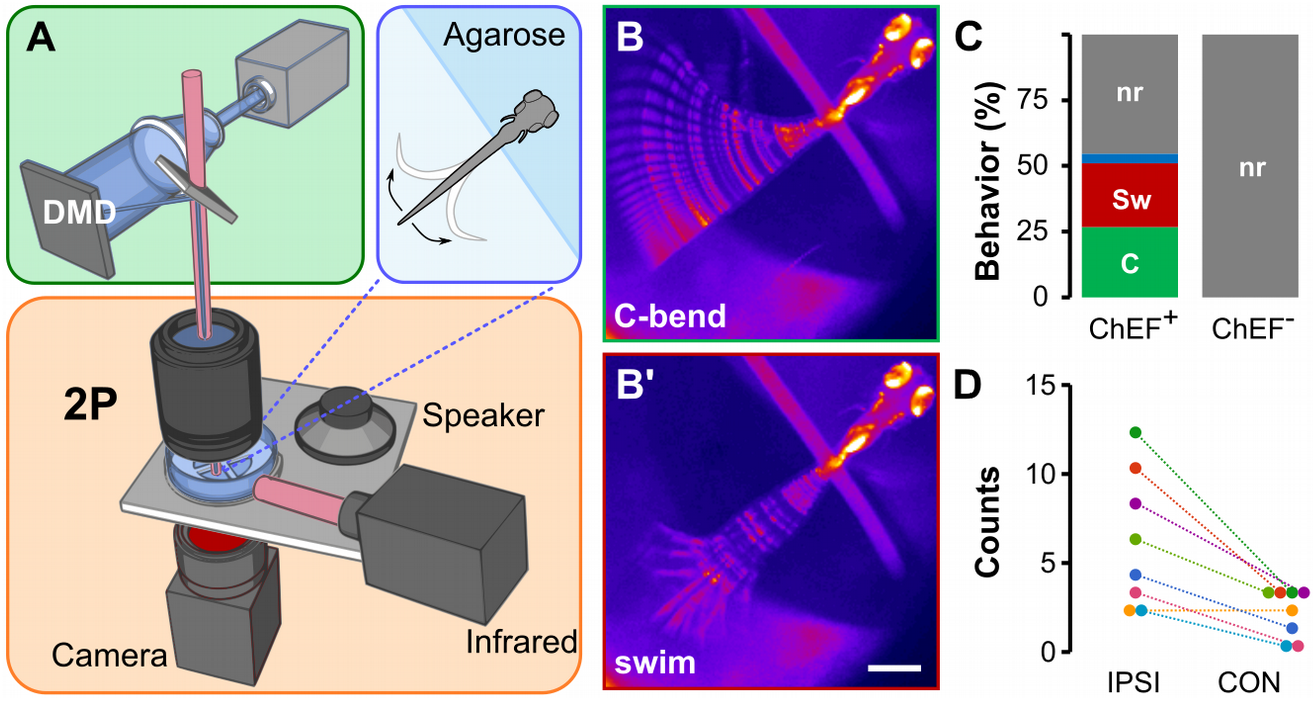
Optogenetic prepontine neuron stimulation elicits C-start behavior (A) Schematic of optogenetic stimulation and two-photon calcium imaging: A digital mirror device (DMD) is used to spatially-restrict 460 nm laser excitation (green box) within the brain of head-embedded larvae (blue box) mounted on a stage with a speaker for acoustic/vibratory stimulation, an infrared light source for tail illumination, and a high-speed camera for behavioral readout (orange box). (B) C-start and swim-like behaviors elicited by unilateral optogenetic stimulation of prepontine neurons in *y293-Gal4*, *UAS:ChEF* positive larvae. Scale bar 500 μm. (C) Percent of behaviors elicited by illumination of larvae expressing ChEF (ChEF^+^; 229 trials, n=8 larvae) and non-expressing sibling controls (ChEF^−^; 63 trials, n=7 larvae). C-start-like responses (C, green), swim-like bouts (Sw, red), other responses (blue), no response (nr, grey). (D) Number of C-start responses made ipsi- and contra-lateral to the side of optogenetic stimulation, color-coded for each of the 8 larvae tested. χ^2^=15.25, * p < 0.001.

Rapid reaction times for Mauthner cell initiated escapes are achieved through a short sensory-motor pathway, use of electrical synapses and the large caliber of the Mauthner axon (Eaton and Hackett, 1984). Thus during fast escapes, the VIIIth nerve directly activates Mauthner neurons, which form mono-synaptic contacts with motor neurons on the contralateral spinal cord (Fetcho, 1992; Yao et al., 2014). As a step toward characterizing the delayed escape pathway, we reconstructed prepontine *y293* neurons. For tracing, we sparsely labeled neurons by crossing *y293-Gal4* to a heatshock-inducible B3 recombinase and a UAS reporter with B3 recombinase ‘blown-out’ (blo) recognition sites (Fig. 4A) (Nern et al., 2011). B3 is relatively inefficient in larval zebrafish, allowing heat-shock conditions to be titrated to achieve stochastic expression of membrane-tagged RFP from the UAS:bloSwitch reporters (Tabor et al., 2018a). We imaged 20 prepontine neurons, and manually reconstructed 5 to visualize their morphology, revealing bilateral terminations in the cerebellar eminentia granularis (EG) and the caudal hindbrain (Fig. 4B, Fig. S5A). A single neurite from each neuron projected ventrally then bifurcated into lateral and medial branches (Fig. 4C). The lateral branch terminated nearby, arborizing in or below the EG (Fig. 4D). The medial branch split again: one fork extended through a dense neuropil area to the caudal hindbrain (Fig. 4E), and the other crossed the midline (through the superior raphe) to the bifurcation zone of neurites from the contralateral prepontine cluster (Fig. 4B-C, arrows). Here, as on the ipsilateral side, the process split, arborizing within the EG (Fig. 4D), and extending to the caudal hindbrain (Fig. 4E). This quadripartite morphology was shared by all 20 neurons imaged, with the only salient differences being (i) the caudal extent of hindbrain projections (Fig. S5B) and (ii) whether neurites projected bilaterally into the EG (Fig. S5C). Caudally projecting neurons did not reach the spinal cord and therefore do not directly activate motor neurons. In addition, neurites were not apposed to VIIIth nerve projections (Fig. S5D-E). Thus, unlike the pathway for fast escapes, where only three synapses are interposed between hair cells and motor neurons, prepontine escape neurons are not directly connected to either sensory input or motor output neurons, which likely contributes to the longer latency of the response.

**Figure 4.**
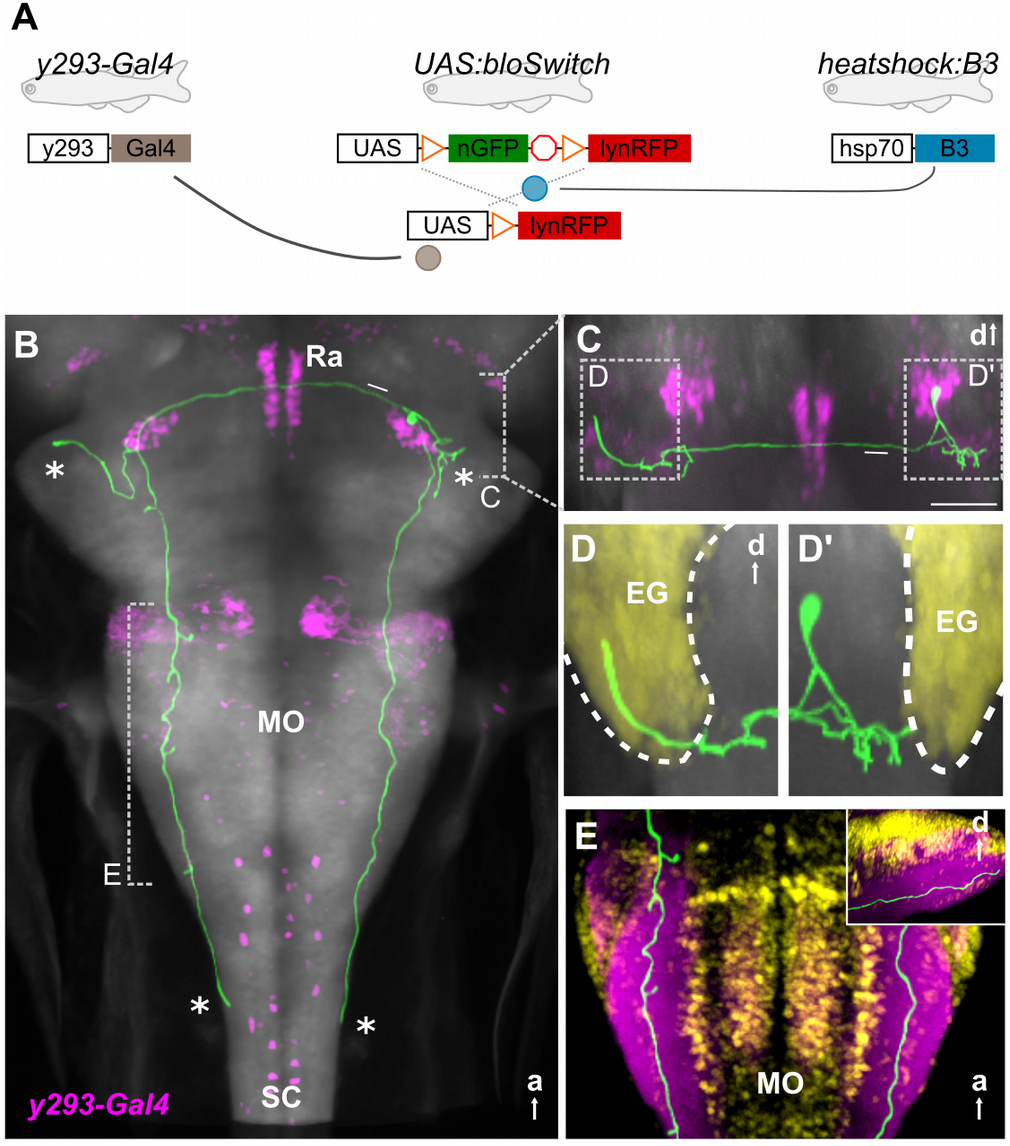
Prepontine escape neurons project reciprocally to the caudal hindbrain and cerebellum (A) Schematic of three transgene system used for B3-recombinase based neuronal tracing. (B-E) Representative traced neuron (green, for others, see Fig. S5), registered to Zebrafish Brain Browser (ZBB) transgene expression atlas (Marquart et al., 2015). Background is *elavl3:Cer* (gray). (B) Horizontal maximum whole-brain projection of a reconstructed neuron from *y293-Gal4* (ZBB, magenta). Asterisks: projections of the four primary neurites. Arrow: commissural projection. Dashed lines indicate views in C and E. a, anterior. (C) Coronal substack projection from the area indicated in B. Arrow: commissural projection. Scale bar 50 um. Views in (D) outlined. a, anterior. (D) Coronal projections of neurites extending into the ipsilateral (D’) and contralateral (D) eminentia granularis (EG, yellow). d, dorsal. (E) Dorsal projection through the caudal medulla lateral neuropil area (ZBB *anti-zrf2*, purple). Cellular regions labeled by *Tg(elavl3:nls-mCar)y517* (ZBB, yellow). Inset: sagittal view of same region. Annotations: EG, eminentia granularis; MO, medulla oblongata; SC, spinal cord; Ra, raphe

We reasoned that the extended pathway and greater reaction time for delayed escapes may provide an opportunity to integrate additional information from the environment to guide LLC trajectories. Because zebrafish larvae are strongly attracted to light, we combined a light spot with an acoustic stimulus and tested escape trajectories (Fig. 5A). In this paradigm, non-directional broad-field illumination on control trials was replaced with a localized light spot several seconds before delivery of a non-directional acoustic stimulus. Whereas SLC trajectories were similar during broad-field illumination and during light-spot exposure, LLC trajectories were preferentially performed toward the spot (Fig. 5B-E). Moreover, body curvature and angular velocity during the initial C-bend were increased during directionalized delayed escapes whereas other kinematic parameters were unchanged (Fig. 5F-G, Fig. S6A). Directionalized responses were absent in *atoh7* mutants, which lack retinofugal projections, confirming that retinal signaling is responsible for guiding delayed escape trajectories (Fig. S6B). Thus, external visual cues strongly influence LLC but not SLC escape trajectory in larvae, consistent with the idea that the longer latency of delayed escapes provides a additional time for integration of sensory information to guide path selection.

**Figure 5.**
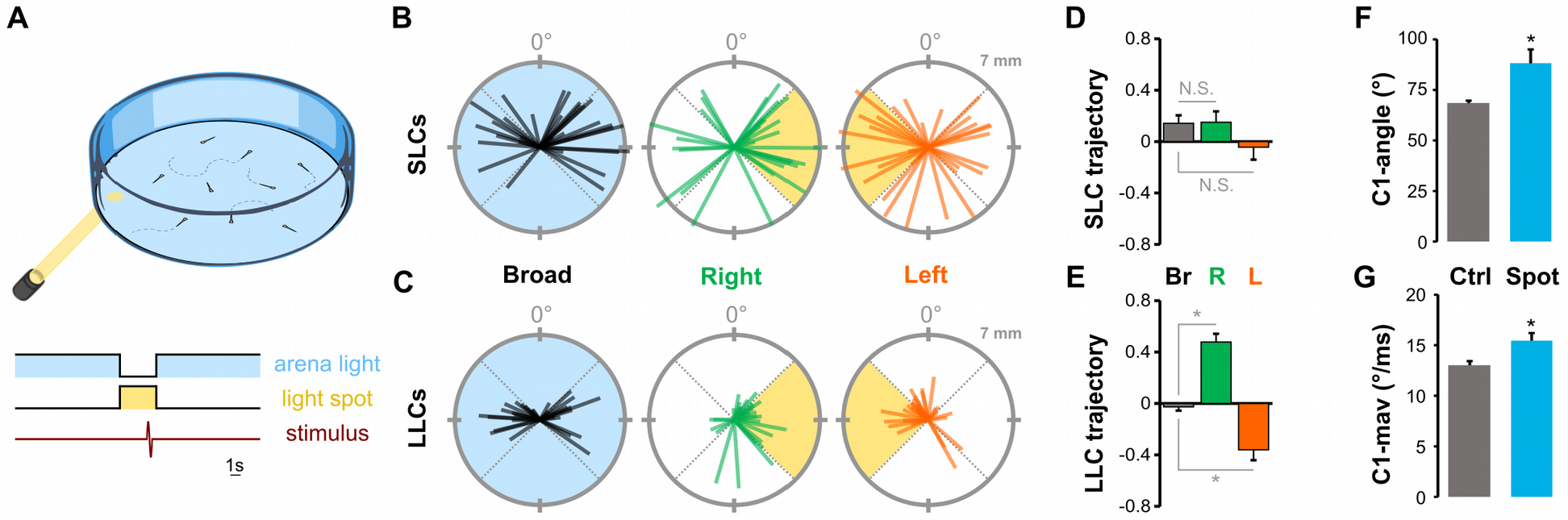
Delayed escape direction is guided by visual information (A) Schematic of experiment measuring escape direction under broad-field illumination or in darkness with only a light spot illuminated. (B-C) 30 representative SLC (B) and LLC (C) escape trajectories of larvae to a non-directional acoustic/vibratory stimulus when under broad-field illumination (Broad), or when oriented to the left or to the right of a light spot. Escape direction is plotted radially and net displacement axially. (D-E) Mean direction choice (−1 all left; +1 all right) for SLC (D) and LLC (E) responses under broad-field illumination (Br; SLC, n=367 responses; LLC, n=372), when the light-spot was to the left of the larva (L; SLC, n=131 responses; LLC, n=257), or to the right of the larva (R; SLC n=143; LLC, n=283). * p < 0.001. (F-G) Mean initial bend angle (F) and maximum angular velocity (G) for LLCs performed under broad-field illumination (grey) or during directionalized responses with a light-spot. * p < 0.01.

To test whether prepontine escape neurons integrate sensory information, we performed two-photon calcium imaging of nuclear-localized GCaMP6s in head-embedded *y293-Gal4* larvae. In parallel, we monitored tail movements in order to correlate activity with behavior. We simultaneously recorded from multiple prepontine *y293-Gal4*, *UAS:GCaMP6s* neurons during presentation of an auditory stimulus, then grouped responses based on the behavioral outcome (Fig. 6A). Mean SLC responsiveness was 20.4 ± 1.5%, however, unexpectedly LLC responses were only elicited on 1.1% of trials in embedded larvae. The low rate of delayed escapes in immobilized larvae precluded us from testing the effect of directionalized light stimuli. Nevertheless, on trials with a delayed escape, prepontine neurons on the same side as the initial C-bend showed a significant increase in mean activity (Fig. 6B-C). Activity was less elevated, but also above baseline, on trials with a delayed escape on the contralateral side. However, neurons were completely inactive on trials where the acoustic stimulus failed to elicit a reaction, demonstrating that the prepontine clusters are not sensory interneurons but motor-associated neurons whose activity correlates most strongly with ipsilateral delayed escape reactions. Strikingly, prepontine escape neurons were also silent on trials where the larva performed a fast escape. This suggests that fast escape and delayed escape pathways can not be co-active and suggests that Mauthner-mediated fast escape responses suppress the delayed escape pathway. We propose that the auditory stimulus recruits independent pathways for escape, and that the faster reaction time of the M-cell pathway shuts down the delayed escape circuit, preventing transmission of potentially conflicting motor commands (Fig. 7).

**Figure 6.**
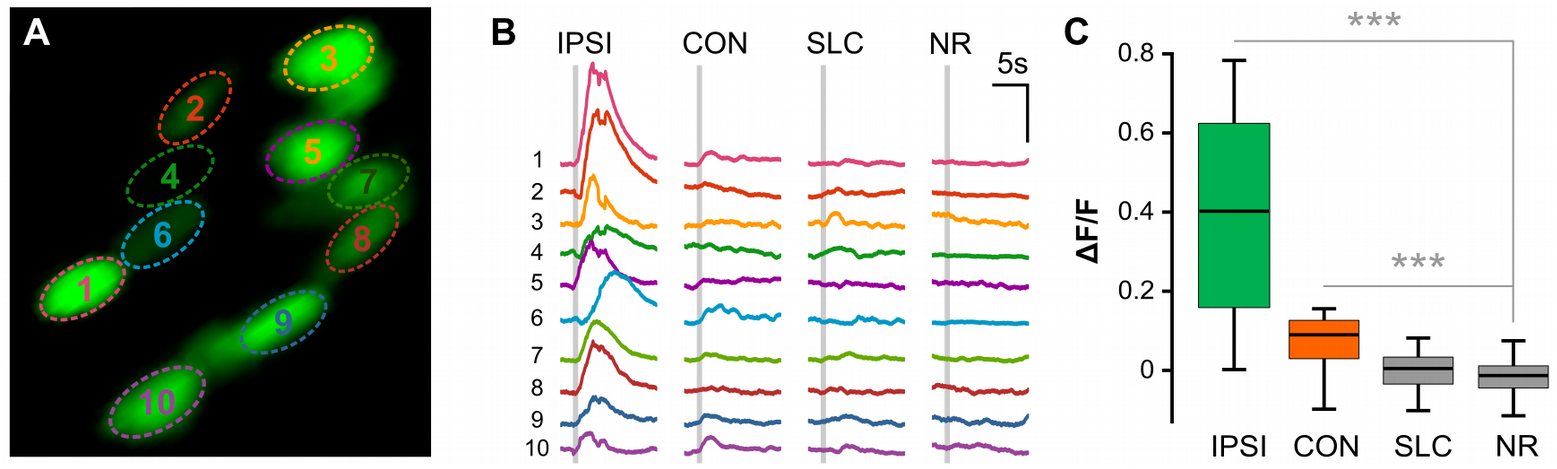
Prepontine neurons are active during ipsilateral delayed escapes (A) Two-photon optical section of nuclear-localized GCaMP6s positive prepontine neurons in *y293-Gal4* with ROIs shown in (B) indicated. (B) Representative GCaMP6s traces for ipsilateral LLCs (IPSI), contralateral LLCs (CON), SLCs, and no response (NR) trials. Grey bar: acoustic stimulus. (C) Change in GCaMP6s fluorescence (ΔF/F) across response types: ipsilateral LLCs (IPSI) contralateral LLCs (CON), SLCs, and no response (NR) trials (31 neurons, n=3 larvae).

**Figure 7.**
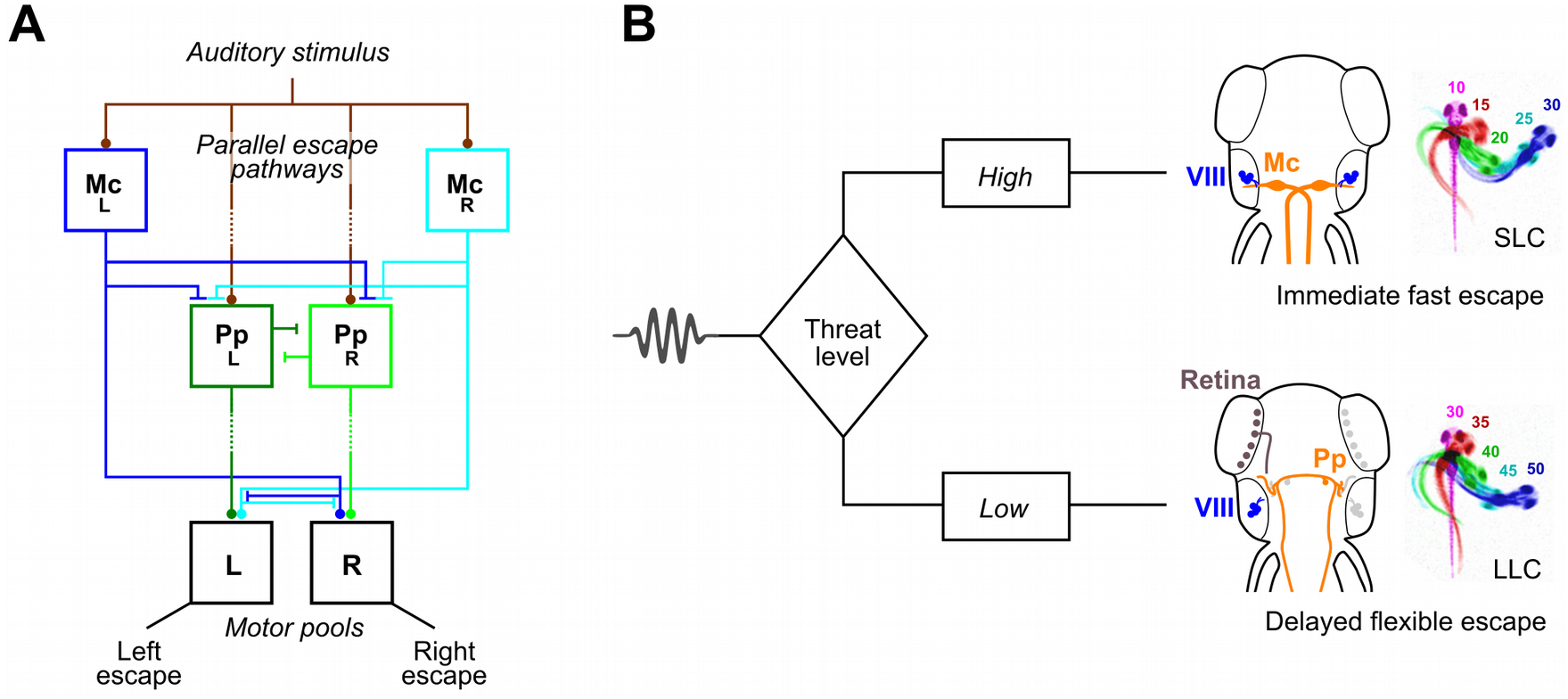
Escape pathways in zebrafish (A) Parallel sensory pathways transmit acoustic information to Mauthner cells (M_L_ and M_R_) and prepontine escape neurons. Decision making is based on reaction time: VIIIth nerve activation of M-cells is direct whereas prepontine neurons receive auditory information only via an indirect pathway allowing active M-cells to prevent the initiation of delayed escapes. (B) Anatomy corresponding to the model in (A). Auditory signals from the statoacoustic ganglion (SAG, brown) excite M-cells (M) directly and prepontine escape neurons (green) indirectly. M-cells receive predominantly ipsilateral inputs and project commissurally to drive fast escapes, whereas prepontine neurons project both ipsi- and contralaterally and may drive escape in either direction.

## Discussion

Rapid escape responses in many species are mediated by giant fiber neurons, providing a conspicuous entry point into the underlying circuit. In contrast, neuronal pathways for alternate modes of escape have not been well characterized. Here, we reveal a population of premotor neurons that initiate delayed escape behavior in larval zebrafish: a bilateral cluster of approximately 20 neurons per side adjacent to the locus coeruleus in the prepontine hindbrain. Although prepontine escape neurons project bilaterally to the caudal medulla oblongata, calcium imaging, lesion and optogenetic activation experiments all indicate that these neurons predominantly initiate ipsilateral escapes. Delayed escape trajectories are strongly biased by visual cues from the environment, suggesting that these responses represent a more ‘deliberative’ mode of escape, potentially allowing larvae to better evade predators or obstacles.

The pathway we describe for auditory induced delayed escapes is not similar to previously described escape circuits in zebrafish. Rapid C-start escapes to head-touch stimuli are mediated by the reticulospinal neurons MiD2cm and MiD3cm (Liu and Fetcho, 1999), and slow velocity looming stimuli trigger non-Mauthner escapes, potentially via a set of reticulospinal neurons that show stimulus-correlated activity (Bhattacharyya et al., 2017). However, unlike reticulospinal neurons, which traverse the medial and lateral longitudinal fasciculi to contact targets in the spinal cord, prepontine escape neurons project through lateral fiber tracts and terminate in a dense neuropil zone in the caudal hindbrain, and must therefore drive spinal cord motor neurons indirectly. Similarly, unlike the Mauthner neurons which receive mono-synaptic auditory input from the VIIIth nerve, prepontine escape neurons must receive polysynaptic inputs, potentially within the eminentia granularis, a region known to receive sensory input and where prepontine neurite morphology resembles dendritic arborizations (Liao and Haehnel, 2012). A third difference is the mechanism for selecting escape direction. Feed-forward inhibitory signals help to select activation of a single M-cell (Koyama et al., 2016); however acoustic stimuli often activate both M-cells and downstream mechanisms prevent simultaneous bilateral activation of motor pools (Satou et al., 2009). In contrast, prepontine neurons show much greater activity on the side ipsilateral to the escape direction. Commissural processes project reciprocally to the contralateral nucleus, raising the possibility that lateral inhibition ensures unilateral activation and initiation of an escape to one side.

Although Mauthner-initiated escape responses in adult fish are biased by visual cues, our data indicated that at larval stages only delayed escape trajectories were biased by visual information (Canfield, 2003). Delayed escapes may provide sufficient time for cross-modal integration and computation of an optimal escape trajectory to evade threats or obstacles. In addition, less vigorous, long-latency escapes may also allow animals to calibrate the cost of behavioral responses to perceived threat (Bhattacharyya et al., 2017). Consistent with this idea, LLCs are preferentially evoked by weak acoustic stimuli, and also match response vigor and speed to stimulus intensity (Burgess and Granato, 2007; Jain et al., 2018). In some circumstances, the predictable path trajectories of fast escapes are susceptible to exploitation (Catania, 2009). Indeed, many prey species show ‘protean behavior’, exhibiting intrinsically erratic or variable responses to confuse predators (Humphries and Driver, 1970). For zebrafish, the presence of alternate modes of escape and the intrinsic variability of delayed escape behavior may reduce the predictability of escape trajectories. These roles are not exclusive: faced with a less precipitous threat, larvae may compute an optimal escape trajectory that is also energetically favorable and more flexible than Mauthner cell driven fast escape reactions.

Delayed escape neurons are located in an unannotated area of the prepontine hindbrain between the locus coeruleus and the cerebellum. The *y293-Gal4* line is an enhancer trap for *fndc5b*, which is expressed in a topographically similar area in mice that is annotated as the vestibular nucleus. This is striking because the VN drives startle responses to abrupt vestibular stimuli in mammals (Bisdorff et al., 1994; Li et al., 2001). The precise pathway for vestibular startle has not been characterized, but is independent of the system that drives startle responses to acoustic or somatosensory stimuli (Steidl et al., 2004). Thus, based on their commissural and cerebellar projections, and location in rhombomere 1 proximal to the LC (Straka et al., 2001), we propose that prepontine escape neurons reside in the zebrafish homolog of the mammalian superior vestibular nucleus, and may represent an evolutionary ancient secondary pathway for rapid defensive responses to threats sensed via acoustic or vibrational cues.

The M-cell system has given us one of the most complete pictures of neural circuit function in vertebrates, however its command-like structure is not representative of how most decisions are computed in vertebrate nervous systems. The identification of neurons that subserve delayed escape reactions now offers the opportunity to study behavioral choice at cellular resolution in an ethologically relevant and experimentally tractable system.

## Materials and Methods

### Animal Husbandry

Gal4 enhancer trap and transgenic lines used in this study were maintained in a Tüpfel long fin (TL) strain background. Embryos were raised in E3 medium supplemented with 1.5 mM HEPES pH 7.3 (E3h) at 28°C on a 14 h:10 h light:dark cycle with medium changes at least every 2 days unless otherwise described. All *in vivo* experimental procedures were conducted according to National Institutes of Health guidelines for animal research and were approved by the NICHD animal care and use committee.

### Mutant and transgenic Lines

Images throughout were registered to the Zebrafish Brain Browser to enable comparison with other markers (Marquart et al., 2015). Gal4 lines used for the circuit-breaking screen were previously described (Bergeron et al., 2012; Marquart et al., 2015), and maintained using *Tg(UAS-E1b:Kaede)s1999t* (*UAS:Kaede*) (Davison et al., 2007). For genetic ablation experiments, lines were crossed to nitroreductase lines *Tg(UAS-E1b:BGi-epNTR-TagRFPT-oPre)y268Tg* or *Tg(UAS:epNTR-TagRFPT-utr.zb3)y362Tg* (Marquart et al., 2015; Tabor et al., 2014). *UAS:bloswitch* and *hsp70l:B3* lines were used for *y293Et* neuron tracing (Tabor et al., 2018a). *Atoh7*^*sa16352*^ mutants were acquired from the Zebrafish International Resource Center (ZIRC)(Busch-Nentwich et al., 2013). The LC was visualized using *Et(gata2a:EGFP)pku2* (*vmat2:GFP*) (Wen et al., 2008). Images of *TgBAC(chata:Gal4-vp16)mpn202* (*chata-Gal4*) were as published (Forster et al., 2017), registered to ZBB.

### Imaging

Embryos were raised in E3h media containing 300 μM N-Phenylthiourea (PTU) starting at 8–22 hpf to suppress melanophore formation with PTU changed at least every 48 hrs. For imaging at 6 dpf, larvae were anesthetized in 0.24 mg/mL tricaine methanesulfonate (MS-222) for 3 min and mounted in 2.5% low melting point agarose in 3D printed plastic inserts (ABS from Stratasys or clear resin from FormLabs) within #1.5 thickness (0.17 ± 0.005 mm) cover glass bottom cell culture chambers (Lab-Tek II 155379). An inverted laser-scanning confocal microscope (Leica TCS SP5 II) equipped with an automated stage and 25x/0.95 NA apochromatic water immersion lens (Leica # 11506340) was used to acquire confocal stacks. For labeling individual neurons, *y293-Gal4; UAS:bloSwitch* fish were crossed to *hsp70l:B3*. Sparse labeling was achieved by 25-35 min heatshock at 37°C at 3 dpf to induce B3 recombinase. Larvae were then raised under standard conditions and imaged at 6 dpf. Neurons were traced in Imaris 8.4.2, exported as TIFs and converted to NIFTI for alignment with ANTs to *y293-Gal4* as a reference (Avants et al., 2008). To photoconvert Kaede from green to red in selected neurons, we scanned with a 405nm laser at 30 mW for 90 s.

### Genetic and laser ablations

For genetic ablations, we used an engineered variant (epNTR) of the bacterial nitroreductase gene, which converts a cell permeable substrate (metronidazole) into a cell impermeable cytotoxin (Pisharath et al., 2007; Tabor et al., 2014). Gal4 enhancer trap lines were crossed to UAS:epNTR and embryos screened for red fluorescence. Non-fluorescent embryos were used as controls. At 4 dpf larvae were exposed to 10 mM metronidazole for 24 hrs, given 24 hrs to recover, and then tested for escape behavior at 6 dpf. For laser excisions, subsets of Kaede-positive neurons were selectively ablated at 4 dpf in *y293-Gal4*;*UAS:Kaede* larvae raised in PTU. Laser excisions were performed on an upright laser-scanning confocal microscope (Leica TCS SP5 II) equipped with a multiphoton laser (SpectraPhysics MaiTai DeepSee), automated stage, and 20x/1.00 NA apochromatic water dipping lens (Leica # 11507701). Larvae to be ablated as well as controls were mounted in 2.5% low melting point agarose on #1.5 thickness cover glass which was then inverted for the upright microscope. A 488 nm argon laser line was used to visualize target cells and confirm ablation, while the multiphoton laser tuned to 800 nm was used for selective laser excision The laser was pulsed for 5-1000 ms at ~2.4W until cell integrity was compromised. Following ablation, larvae were raised in E3h until behavioral testing at 6 dpf. Successful ablations were then confirmed by confocal microscopy — only larvae with 3 cells or less remaining on either side were analyzed.

### Free-swimming behavior

For light spot experiments, TL larvae were tested in groups of 15-20 within a 33 × 33 mm corral, which kept larvae in view of a high-speed camera. Larvae were illuminated from above at ~80 μW/cm^2^ (arena light) and by an infrared LED array for imaging purposes from below. During light spot trials, the arena light was replaced for 3.5 sec with a light spot of ~8 μW/cm^2^ and ~6 mm diameter focused from below. 3 sec after appearance of the light spot, larvae were exposed to an acoustic/vibratory stimulus. Aside from the switching of illumination, control trials to quantify baseline responses were performed under the same conditions. Control and light spot trials were pseudo-randomly presented 20-40 times each at 15 sec intervals. The change in illumination elicited O-bends only within the first second, and not at 3 sec when the acoustic stimulus was provided. For genetic ablation experiments, larvae were tested individually in 9.7 × 9.7 mm wells of a 3×3 grid illuminated by an LED array at ~500 μW/cm^2^ from below. Different stimulus intensities were presented 20 times each in a pseudorandom sequence at 15 sec intervals to minimize habituation. Genetically ablated larvae and metronidazole-treated controls were tested in alternation.

Auditory stimuli consisted of sinusoidal waveforms of 21 to 36 dB, of 2 ms duration, and nominally 250 or 1000 Hz, though the acoustic/vibrational stimuli as delivered are intrinsically broadband. Stimuli were delivered with an electrodynamic exciter (Type 4810 Mini-shaker; Brüel & Kjær) controlled by an digital–analog data acquisition card (PCI-6221; National Instruments). Behavioral responses of larvae were recorded at 1,000 frames/s with a high-speed camera (DRS Lightning RDT/1 or RL Redlake MotionPro; DEL Imaging) fitted with a 50 mm macro lens (EX DG Macro, Sigma Co.). For light spot experiments requiring infrared illumination, cameras were additionally fitted with an infrared filter (R72, Hoya Filters). Recorded trial bouts were 120 ms in length with acoustic/vibratory stimuli delivered at 30ms, except for laser excisions of *y293-Gal4* larvae where we analyzed 200 ms to ensure that LLCs were absent and not merely delayed.

Behavioral responses were analyzed with Flote software (Burgess and Granato, 2007). As ongoing locomotion differentially influences SLC and LLC probability, larvae were only included if they were motionless in the 30 ms prior to acoustic stimulus. Startle responses were considered SLCs if they occurred within 12 ms of stimulus delivery. LLC responsiveness was calculated as the mean proportion of larvae responding with a long-latency C-starts, as a fraction of all larvae still stationary after the time period during which SLC responses occur (Burgess and Granato, 2007). This adjustment is made because SLC production precludes the production of LLCs. For light spot experiments, larvae were pooled by quadrant based on their initial orientation to the light spot. For behavioral analysis following PTU treatment, some low-contrast larvae were unable to be automatically tracked with Flote and behaviors were manually assessed with the scorer blinded to the identity of ablated versus control conditions.

### Calcium imaging and optogenetic activation

For calcium imaging or optogenetic stimulation, *y293-Gal4* embryos were injected at the one cell stage with tol1 mRNA and a plasmid containing either *UAS:BGi-nls-GCaMP6s.zf2-v2a-nls-dsRed.zf1-afp* or *UAS:BGi-ChEF-v2a-mCherry-afp* respectively (Tabor et al., 2018a), screened for red fluorescence and raised in PTU in the dark. At 6 dpf, GCaMP6s- or ChEF-positive larvae were embedded in 3.5% low melting point agarose in E3h in a Petri dish with agarose once hardened cut away from the tail caudal to the swim bladder to allow for tail movement and behavioral readout of acoustic or optogenetic stimulation. Larvae were then place on a custom 3D printed stage with temperature maintained at 28 °C by a ring-shaped Peltier device. To track tail movements, larvae were illuminated using an 980 nm LED and imaged from below at 100 or 200 frames per second using an infrared CCD camera (Pike F-032C IRF, Allied Vision Technologies). Tail movements were acquired and tracked using custom Matlab script. Each larva was tested with a mean of 8 trials at 5-10 minute intervals. A trial comprised 4 stimuli at an interstimulus intervals of 60-90 s. Only larvae that performed both SLCs and LLCs were included for analysis.

GCaMP6s- or ChEF-positive neurons were imaged on a custom-built multiphoton microscope with a 20x/0.90 NA water dipping lens (Olympus) and a Ti-Sapphire laser (Coherent Chameleon Vision-S) tuned to 950 nm for excitation and controlled in Matlab (Mathworks) by ScanImage (Pologruto et al., 2003). For calcium imaging, GCaMP6s signals from single planes through left or right prepontine neurons were imaged at 1.95-13.95 fps. For optogenetic stimulation, captured images were converted into binary ROIs and projected back onto the larval zebrafish brain by a digital micromirror device (DLi4130, Digital Light Innovations) for durations of 10 or 100 ms controlled by Clampex (pCLAMP 10.4, Molecular Devices). GCaMP6s ΔF/F was quantified in nuclear ROIs drawn in Fiji with the frames representing ~1 sec averaged and compared 2.72 sec after acoustic stimulation compared to those within 1 sec immediately prior. Trials with spontaneous tail movement within 100 ms prior to acoustic stimulation were excluded from analysis.

### Statistics

Analysis was performed with IDL (Harris), R (http://www.R-project.org/) and Gnumeric (http://projects.gnome.org/gnumeric/). Graphs and text report means and standard errors. Box plots show median and quartiles; whiskers show 10-90%. Bar plots show mean and standard error. N reported in figure legends.

## Acknowledgements

This work was supported by the Intramural Research Program of the *Eunice Kennedy Shriver* National Institute for Child Health and Human Development (NICHD) and utilized the high-performance computational capabilities of the Biowulf Linux cluster at the National Institutes of Health, Bethesda, MD.

---

Figure S1. Additional motor phenotypes after chemogenetic neuronal ablation

Kinematic measures for LLC (A) and SLC (B) responses after ablating neurons labeled in *y252*, *y293* and *y330*

Figure S2. Prepontine escape neurons are located between the locus coeruleus and cerebellum

(A-B) Dorsal (A) and parasagittal (B) projections from ZBB of *y293-Gal4*, *y264-Gal4*, and *chata-Gal4* labeled neurons in rhombomeres 1-4 (R1-4). Prepontine neurons labeled by *y293-Gal4* are located in rhombomere 1, in contrast to the anterior and posterior trigeminal motor nuclei labeled by *chata-Gal4* located in R2 and R3 respectively and the Mauthner-cell in R4 labeled by *y264-Gal4*.

(C) Coronal projection of *y293-Gal4* prepontine neurons situated between the locus coeruleus (LC) and the cerebellum (CE).

Figure S3. Additional phenotypes after laser ablation of neurons in rhombomeres 1 and 6

(A-B) SLC (A) and LLC (B) responsiveness after R1 ablation (N=9, green), R6 ablation (N=14, blue) and unablated sibling controls (N=27, black). Significant effects of R1 ablations on LLC and SLC probability; ANOVA F_1,102_=23.37, p < 0.001 and F_1,102_=21.79, p < 0.001 respectively.

(C) SLC kinematic measurements after bilateral R1 laser ablation (N’s as above). No significant differences.

(D) SLC directionality (%Right: percent of SLC responses initiated to the right) after unilateral (left) R1 laser ablation. N=13 (ablated) and 24 (control).

Figure S4. Prepontine escape neurons in fish are similar to the mouse superior vestibular nucleus

(A) Bottom: Coronal projection through zebrafish rhombomere 1 (slice 512-517 from ZBB) with nuclear labeling on the left (*elavl3:nls-RFP* in purple) and neuroanatomic segmentation on the right (magenta, optic tectum; yellow, cerebellum; pink, medulla oblongata; gray, neuropil). Top: Coronal projection of the outlined region showing *y293-Gal4* (green, prepontine neurons), *neurod:GFP* (yellow, cerebellum)(Obholzer et al., 2008) and *y405-Gal4* (magenta, locus coeruleus). *y405-Gal4* is an enhancer trap for *roundabout guidance receptor 2* (*robo2*) with strong expression in the locus coeruleus (Tabor et al., 2018b).

(B) Bottom: Mouse post-natal day 56 (P56) coronal section (slice 111 from the Allen Mouse Brain Atlas, AMBA) with Nissl-staining on the left (purple) and neuroanatomic segmentation on the right (magenta, superior colliculus; yellow, cerebellum; pink, medulla oblongata; gray, fiber tracts). Top: AMBA in situ hybridization images for *Robo2* (B) and *Fndc5* (B’). Image credit: Allen Institute, modified from Allen Developing Mouse Brain Atlas.

CE, cerebellum; HB, hindbrain; LAV, lateral vestibular nucleus; LC, locus coeruleus; MV, medial vestibular nucleus; OT, optic tectum; Pp, prepontine escape neurons; SUV, superior vestibular nucleus.

Figure S5. Traces of individual prepontine escape neurons

(A) Dorsal standard deviation projections of five traced prepontine escape neurons with *elavl3:Cer* as a reference (grey).

(A’) Overlay of co-registered neurons in A showing conserved quadripartite morphology. Dotted lines indicated areas expanded in (B) and (C).

(B) Enlargements of hindbrain from A’ with arrowheads indicating neuron terminals.

(C) Enlargements of lateral rhombomere 1 from A’ with arrowheads termini in the cerebellar eminentia granularis (EG).

(D) Horizontal projection of confocal stack including neuron cell bodies (*y293-Gal4; UAS:KaedeR*, red) after selective photoconversion of Kaede to red and statoacoustic ganglion axon rostral termini (SAG, *y256-Gal4;UAS:KaedeG*, green).

(E) Projection of a reconstructed neuron (green) registered to the ZBB, with the *y256-Gal4* pattern (magenta) that labels the SAG.

Figure S6. Movement kinematic measures for LLCs directionalized by a light spot

(A) Kinematic parameters for acoustically-evoked LLCs performed under broad-field illumination (black) or in the presence of a light spot (blue). C1, initial C-start bend. C2, counterbend. * p < 0.01, n=29 groups of larvae.

(B) Percent of LLCs in a right-ward direction in the presence of a light spot for *atoh7* mutant larvae and siblings. *** p < 0.001, * p < 0.05, n=5 plates each atoh7^−/−^ and siblings.

Video S1. Optogenetic stimulation of *y293-Gal4* prepontine neurons

Representative optogenetic trials from three ChEF positive and a ChEF negative control larvae (bottom right) showing behavioral results to patterned illumination and optogenetic stimulation of prepontine neurons in *y293-Gal4*. 460 nm stimulation for 10 or 100 ms is indicated by a red square in the top right corner of each sub-frame with timestamp at the bottom right.

