## Supplementary figures and images for "Prepontine non-giant neurons drive flexible escape behavior in zebrafish"

### Supplemental Figures 1-6

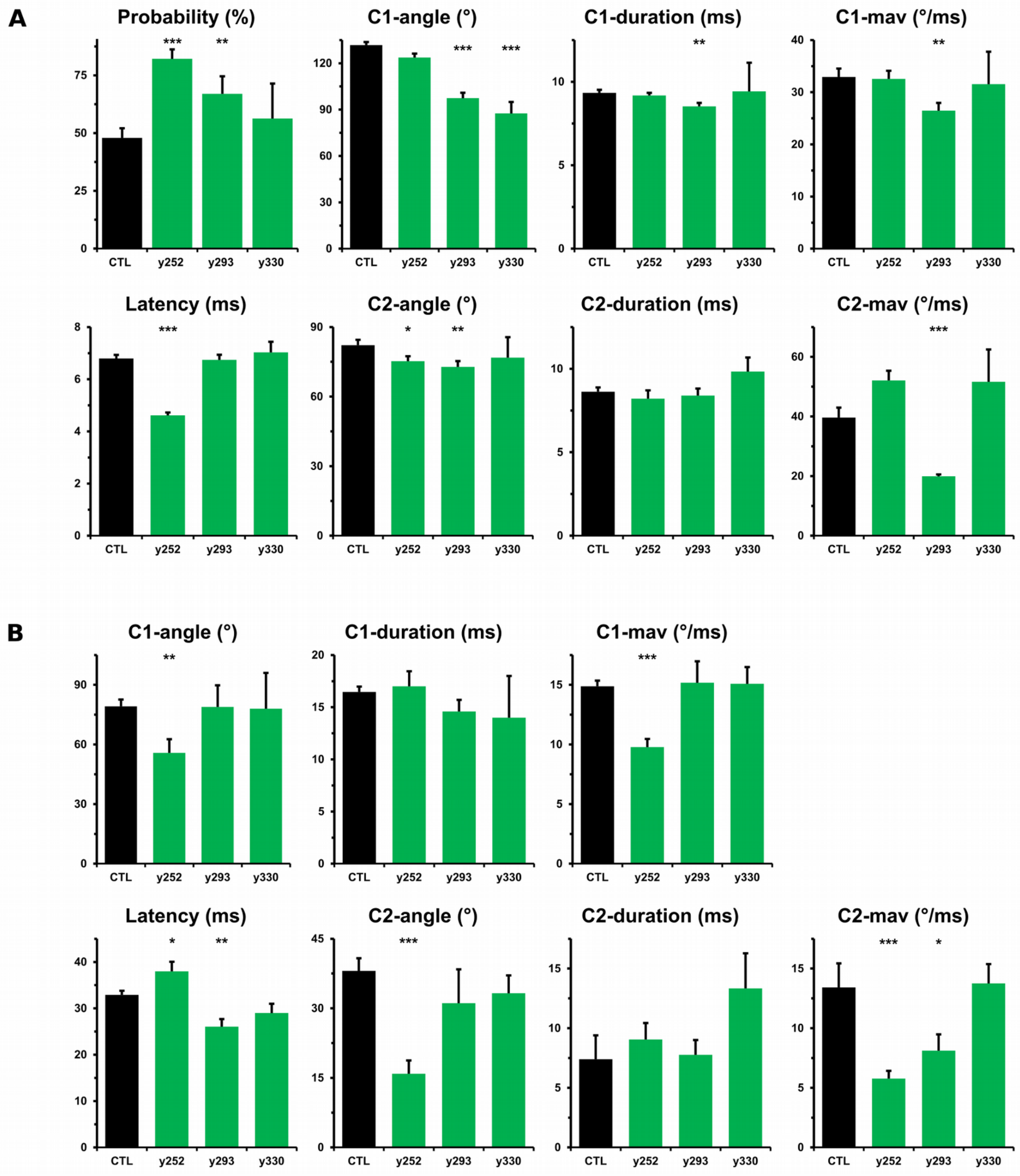

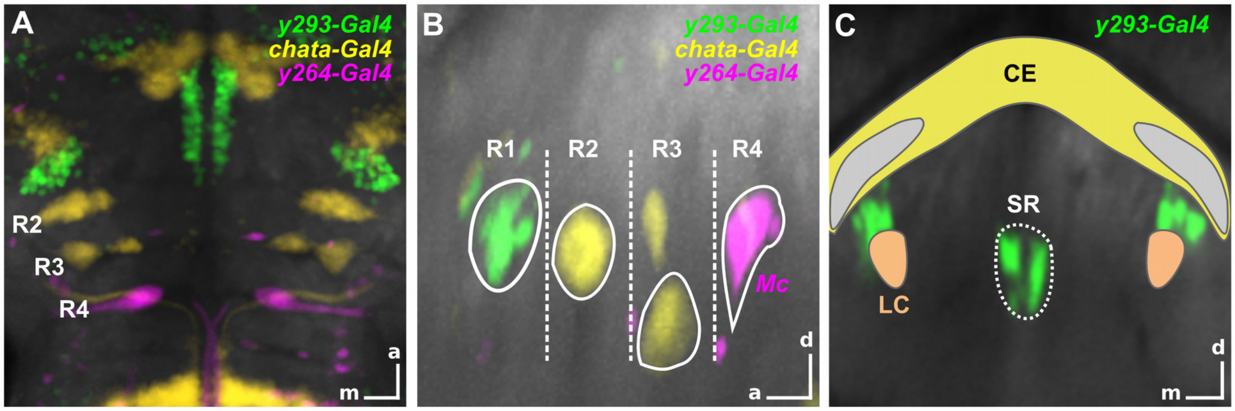

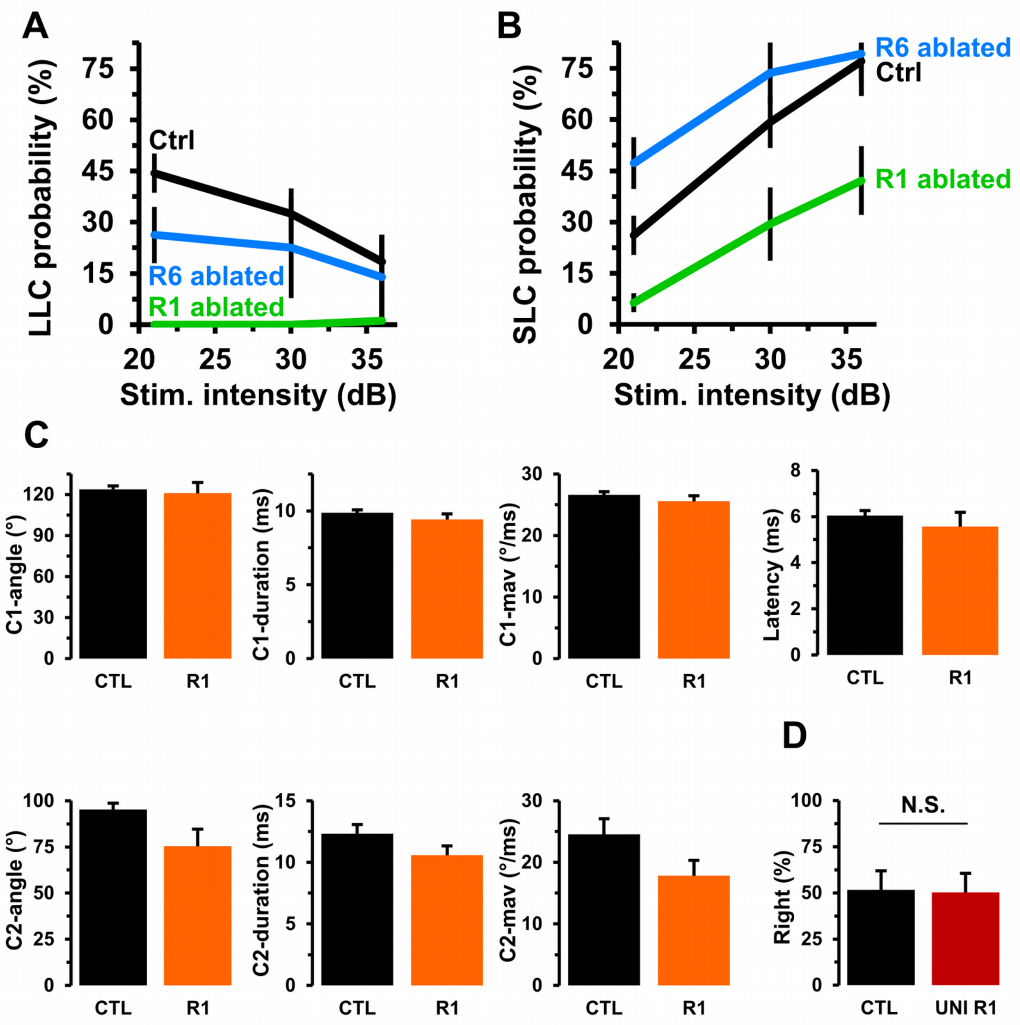

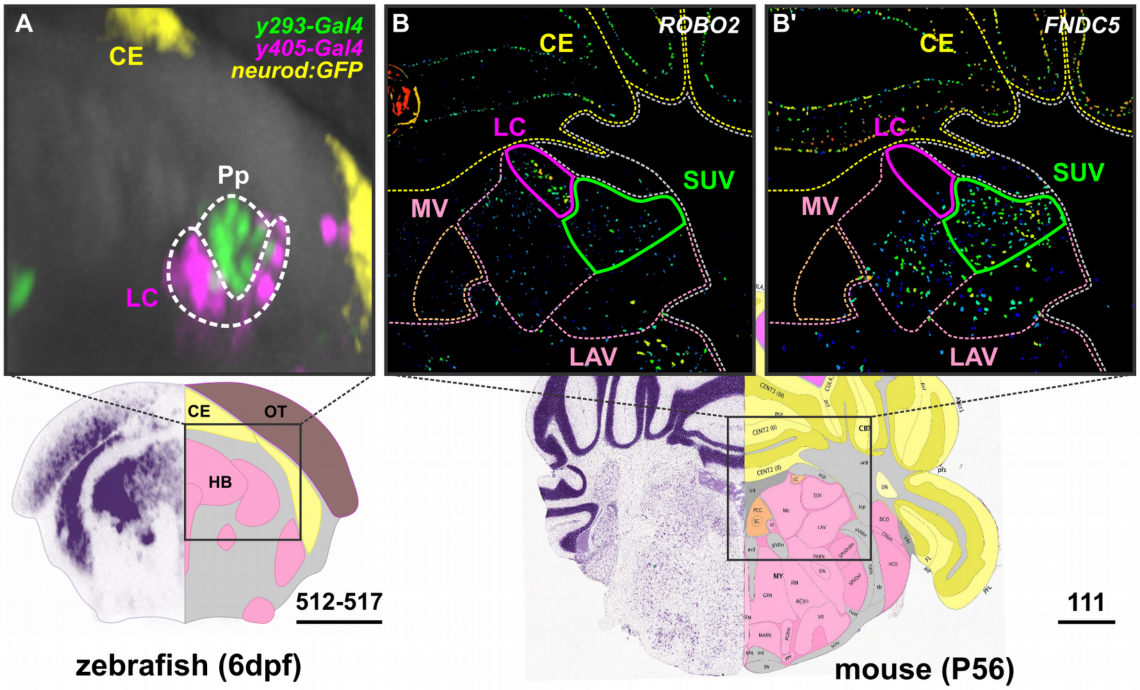

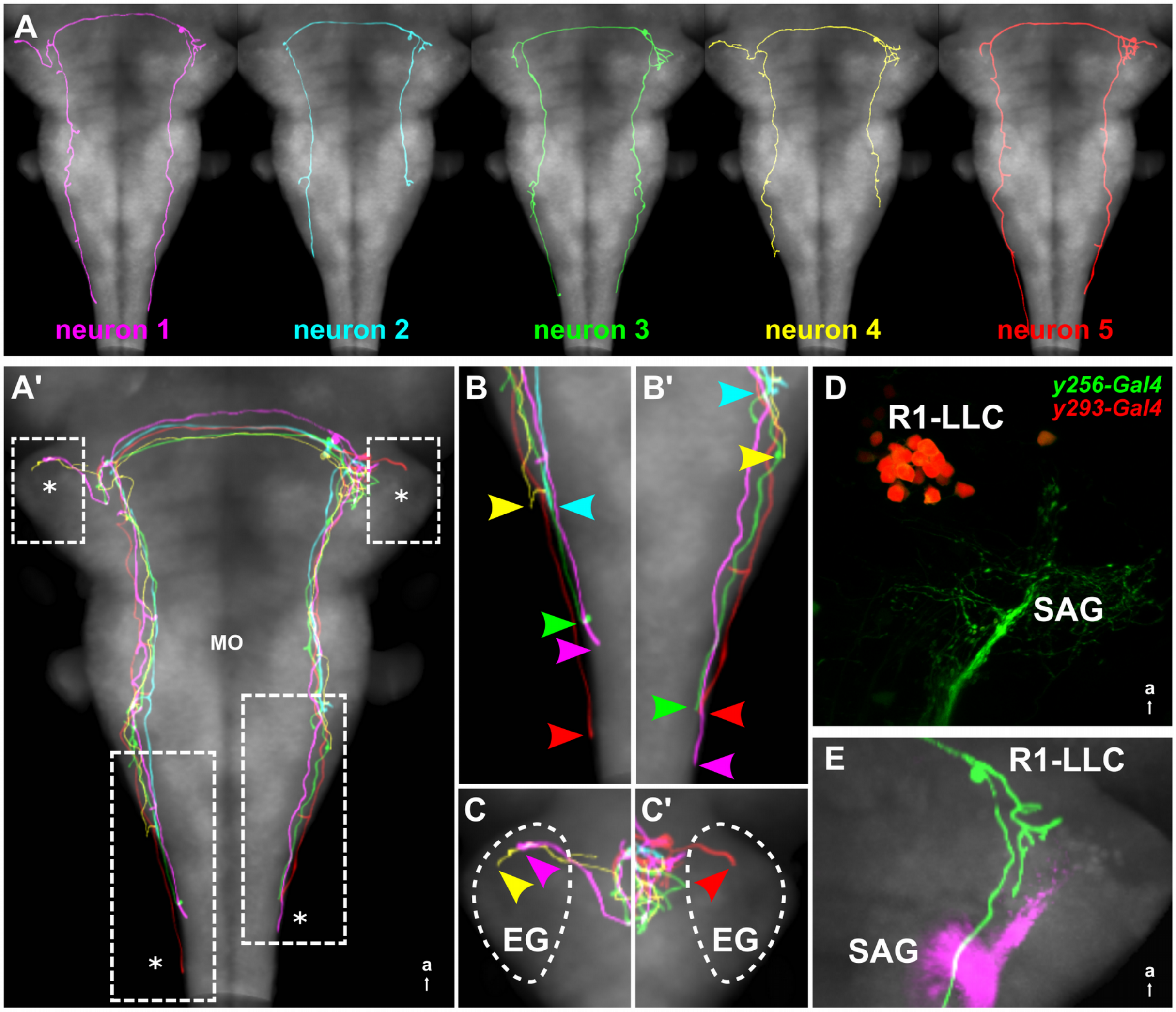

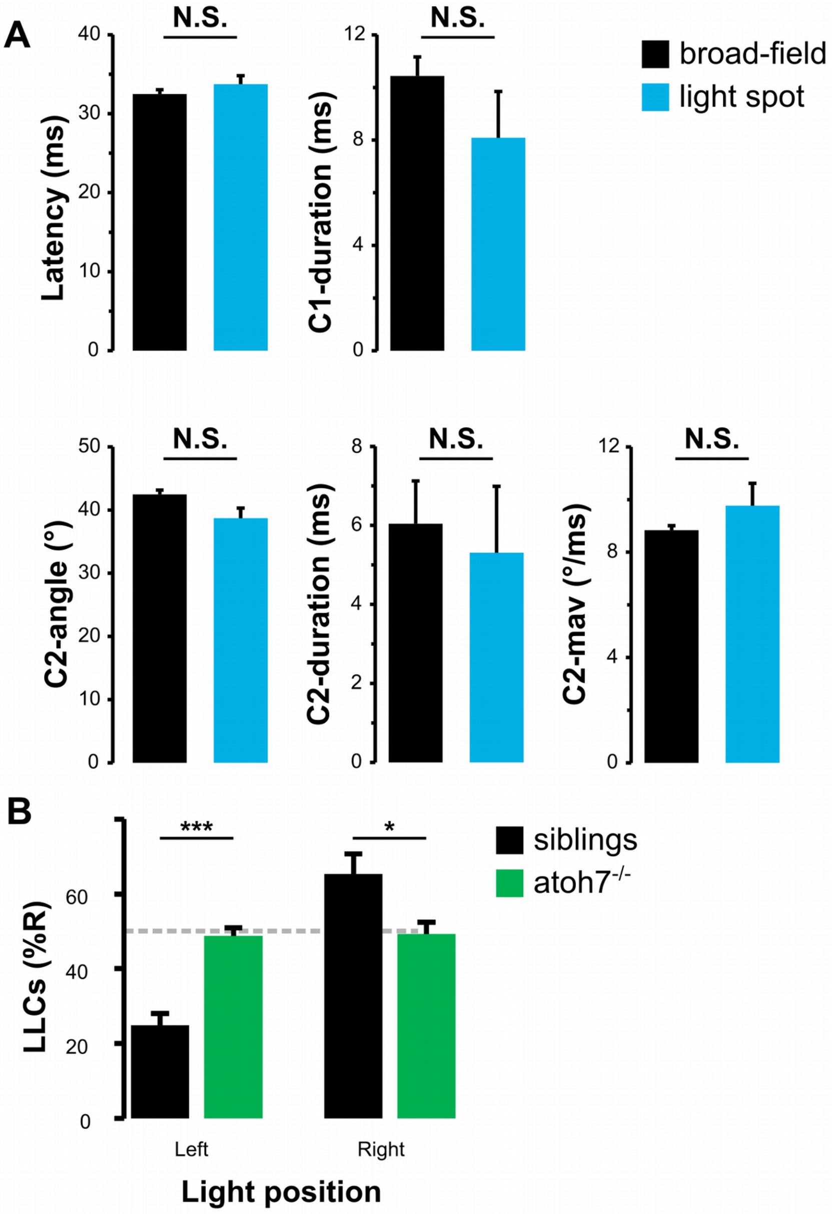
